# Feasibility of an Inhalable Ultrasound Contrast Agent to Enhance Airway Imaging

**DOI:** 10.1101/2021.05.27.446053

**Authors:** Phillip W. Clapp, Phillip G. Durham, Jamie C. Antinori, Rachel W. Walmer, Jessica G. Chlebowski, Brian Velasco, Samantha J. Snow, Paul A. Dayton, Melissa C. Caughey

**Author notes:** **Corresponding Author** Melissa C. Caughey, Joint Department of Biomedical Engineering, University of North Carolina and North Carolina State University, Chapel Hill, NC 27599.

## Abstract

**Introduction:** Ultrasound is a relatively inexpensive and non-ionizing imaging modality, but is under-utilized in large airway assessments due to poor image quality. No commercially available contrast agents currently exist for sonographic evaluation of the respiratory system, nor has a respiratory route of microbubble contrast agent (MCA) administration been previously described for the enhancement of airway imaging.

**Methods:** We conducted a feasibility study to assess proof-of-concept for an inhalable ultrasound MCA composed of lipid-encapsulated decaflourobutane gas. The MCA was nebulized and administered as an aerosol through the lumen of an *ex vivo* porcine trachea, with image enhancement evaluated by comparing images pre- and post-exposure. Additionally, primary human bronchial epithelial (hBE) cells from three donors were differentiated at an air-liquid interface and exposed apically to 25 μL of undiluted MCA or vehicle control to assess contrast agent-induced cytotoxicity and inflammation. Basolateral medium was collected 24-hours post-exposure and lactate dehydrogenase (LDH) and interleukin-8 (IL-8) concentrations were measured as biomarkers of cytotoxicity and inflammation, respectively.

**Results:** Contrast microbubbles remained intact following nebulization and enhanced sonographic delineation of *ex vivo* porcine tracheal walls, indicating adherence of the nebulized MCA to the lumenal mucosa. No significant cytotoxic or inflammatory effects were observed in cultured hBE cells following exposure to MCA.

**Conclusions:** We present proof-of-concept for an inhaled MCA for the enhancement of sonographic evaluations of the large airways. Pending further evaluations for safety and effectiveness, inhaled MCA may be feasible for clinical ultrasound applications, such as enhancing ultrasound-guided tracheal intubation, detecting airway bleeds, or monitoring large airway diseases in pediatric populations.

## Introduction

Multidetector computed tomography (MDCT) is the primary imaging modality used to monitor pathologies of the large airways (>2 mm in diameter),^1^ which include bronchial wall thickening, bronchiectasis (dilation), sacculations, abscesses, nodules, and obstructive mucus impaction.^2^ Although MDCT has excellent resolution, it also has significant drawbacks; namely, exposing the patient to ionizing radiation, time delays in image acquisition and processing, and high cost. Unlike MDCT, ultrasound is inexpensive, non-ionizing, portable, and suitable for point-of-care assessments. However, sonographic evaluations of the airways are challenged by poor ultrasound beam transmission through the air spaces, resulting in suboptimal image quality.

Intravenously administered microbubble contrast agents (MCAs) are U.S. Food and Drug Administration (FDA) approved for the enhancement of cardiac and hepatic ultrasound images, and have a lower risk profile than iodinated contrast agents used to enhance MDCT.^3,4^ MCAs are composed of lipid or albumin outer shells filled with an inert, high molecular weight gas such as sulfur hexafluoride or octafluoropropane (both of which are FDA approved),^4^ or alternatively, decafluorobutane (pending FDA approval). The gas core of MCAs is highly compressible and oscillates when exposed to an ultrasound pulse, causing contraction or expansion of the microbubble diameter (in the range of 1-5 microns) and effective scattering of the ultrasound beam. However, there are no commercially available MCAs for sonographic evaluation of the respiratory system, nor has a respiratory route of MCA administration been previously described for the enhancement of airway imaging. In this report, we describe the feasibility of an inhalable MCA to optimize sonographic delineation of the large airways.

## Methods and Results

We conducted a proof-of-concept study to assess the feasibility of a respiratory route of MCA administration to enhance airway imaging. As described below, sonographic image enhancement was evaluated by administering aerosolized MCA through a three-inch segment of *ex vivo* porcine trachea. Cytotoxic and inflammatory effects of MCA were analyzed in well-differentiated primary human bronchial epithelial (hBE) cells cultured at an air-liquid interface.

### Ultrasound equipment

Imaging was performed using a Siemens Acuson Sequoia ultrasound machine (Siemens Healthineers, Seattle, WA) interfaced with a 15-8 MHz linear probe. The contrast pulse sequencing mode was selected to optimize signal enhancement, setting the mechanical index to 0.19 to prevent destruction of the contrast microbubbles by acoustic energy.^5^

### Nebulized microbubble contrast agent

Microbubbles were formulated with a phospholipid outer shell of distearoylphospatidylcholine (DSPC) and distearoyl-sn-glycero-3-phosphoethanolamine-N-[amino(polyethylene glycol)-2000] (DSPE-PEG2000) and filled with inert, high molecular weight decafluorobutane gas. The microbubbles were activated by mechanical agitation for 45 seconds. Aerosols were generated from the activated MCA using a modified Pari LL nebulizer and Vios compressor (PARI; Starnberg, Germany).

To ensure presence of intact microbubbles following the nebulization process, we aerated a 250 mL beaker of water for 10 seconds with nebulized MCA, to compare with 250 mL of water aerated for 10 seconds by nebulized saline. The submerged microbubbles were imaged continuously from the time of aeration until 30 seconds post-aeration, with the ultrasound transducer held just below the surface of the water. Aeration by nebulized saline resulted in modest bubble formation and echo brightening, which quickly dissipated by 30 seconds (**Figure 1A**). In contrast, aerating the water with nebulized MCA generated strong and persistent echo brightening observed 30 seconds post-aeration (**Figure 1B**).

**Figure 1:**
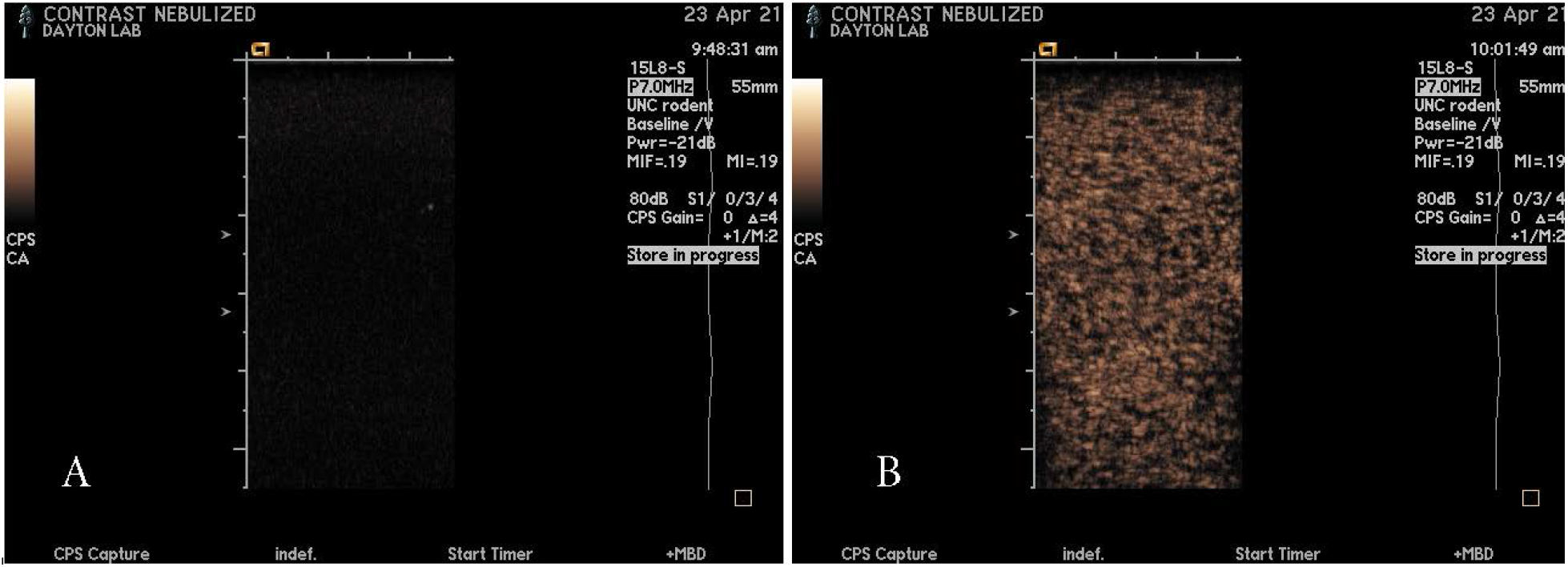
Visualization of water-submerged microbubbles 30 seconds after aerating a water beaker with nebulized saline (Panel A) and nebulized ultrasound contrast agent (Panel B).

### Image enhancement of an ex vivo porcine trachea

To assess image enhancement by nebulized MCA, we constructed a phantom by encasing a fresh three-inch segment of *ex vivo* porcine trachea in a 3.5% agar gel (**Figure 2**). The *ex vivo* trachea was harvested from a 70 kg pig and placed in Ringers solution on ice. A cylindrical cast was used to set the agar, with the *ex vivo* trachea placed 2 cm from the cast surface. Tracheal imaging was performed in the transverse orientation, before and after administration of nebulized MCA. There was no discernable delineation of the tracheal contours prior to ventilation by nebulized MCA (**Figure 3A**). Within 30 seconds of MCA administration, the contour of the near wall of the trachea was easily visualized (**Figure 3B**), demonstrating adhesion of the nebulized MCA to the luminal mucosa. There was no opacification of the air-filled lumen by nebulized MCA, which is consistent with poor ultrasound beam transmission through the air column. By the same principle, there was substantial image dropout beyond the air column, with no visualization of the far wall of the trachea. However, the complete cross-sectional structure of the trachea was easily visualized by obtaining images from the frontal, right lateral, and left lateral imaging planes. To confirm signal enhancement by the nebulized contrast agent, the mechanical index was subsequently increased, which induced microbubble destruction and loss of visualization.

**Figure 2:**
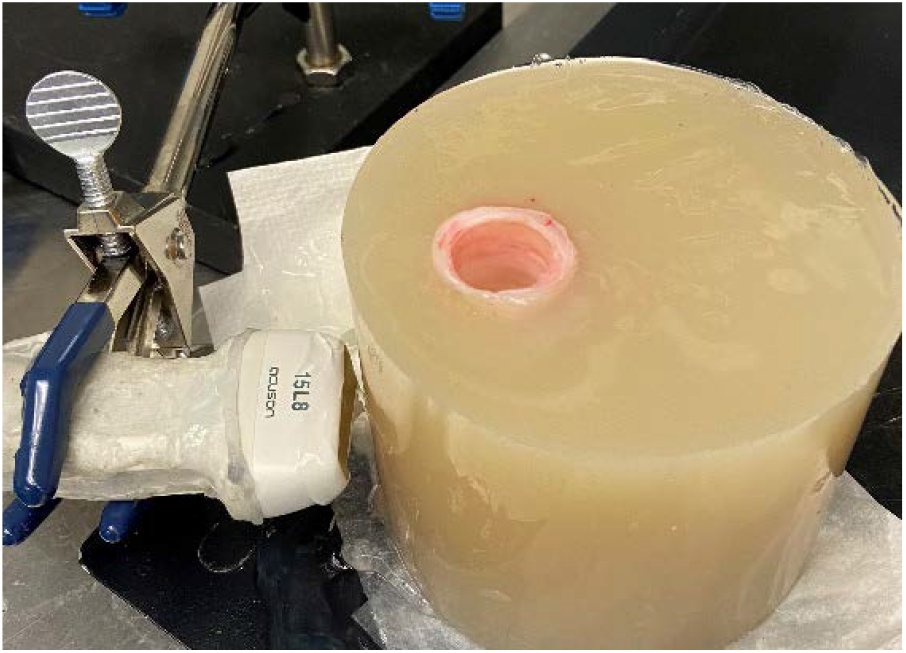
Imaging phantom constructed from *ex vivo* porcine trachea encased in agar.

**Figure 3:**
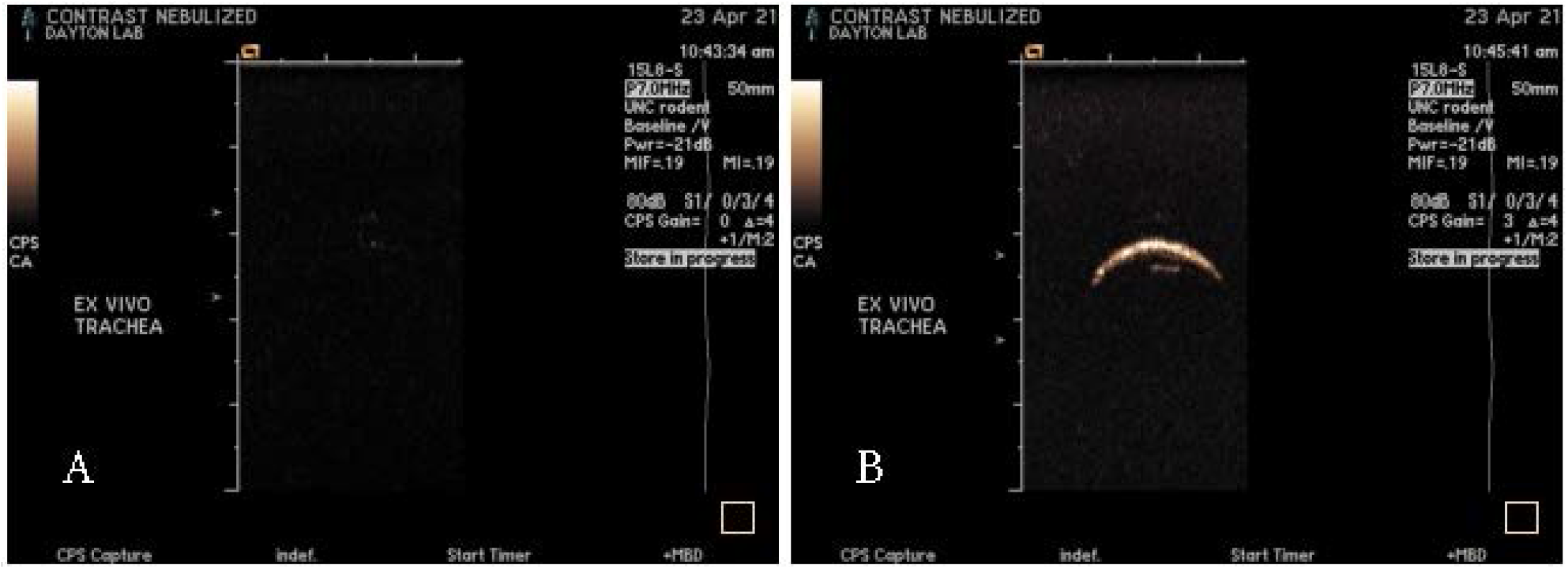
Visualization of *ex vivo* tracheal wall before (Panel A) and after (Panel B) ventilating nebulized microbubble contrast agent (MCA) through the tracheal lumen. Images acquired in contrast pulse sequencing (CPS) mode using an Acuson Sequoia ultrasound system

### Cytotoxicity of MCA to cultured primary hBE cells

We assessed the toxicity of MCA using primary hBE cells collected from 3 donor lungs and differentiated at an air-liquid interface on 12 mm Transwell permeable support membranes to a mucociliary phenotype. Lactate dehydrogenase (LDH) was quantified in the basolateral medium following a 24-hour apical exposure to either 25 μL of undiluted MCA (with the microbubbles activated by mechanical agitation for 45 seconds), 25 μL of undiluted vehicle control, or a no treatment control. One Transwell from each donor was treated with 4% Triton X-100 to lyse cells and provide a positive (maximal LDH release) control. Each exposure condition was normalized to the basolateral LDH content in the positive control condition. Neither the activated MCA nor vehicle control produced cytotoxic responses (**Figure 4**).

**Figure 4:**
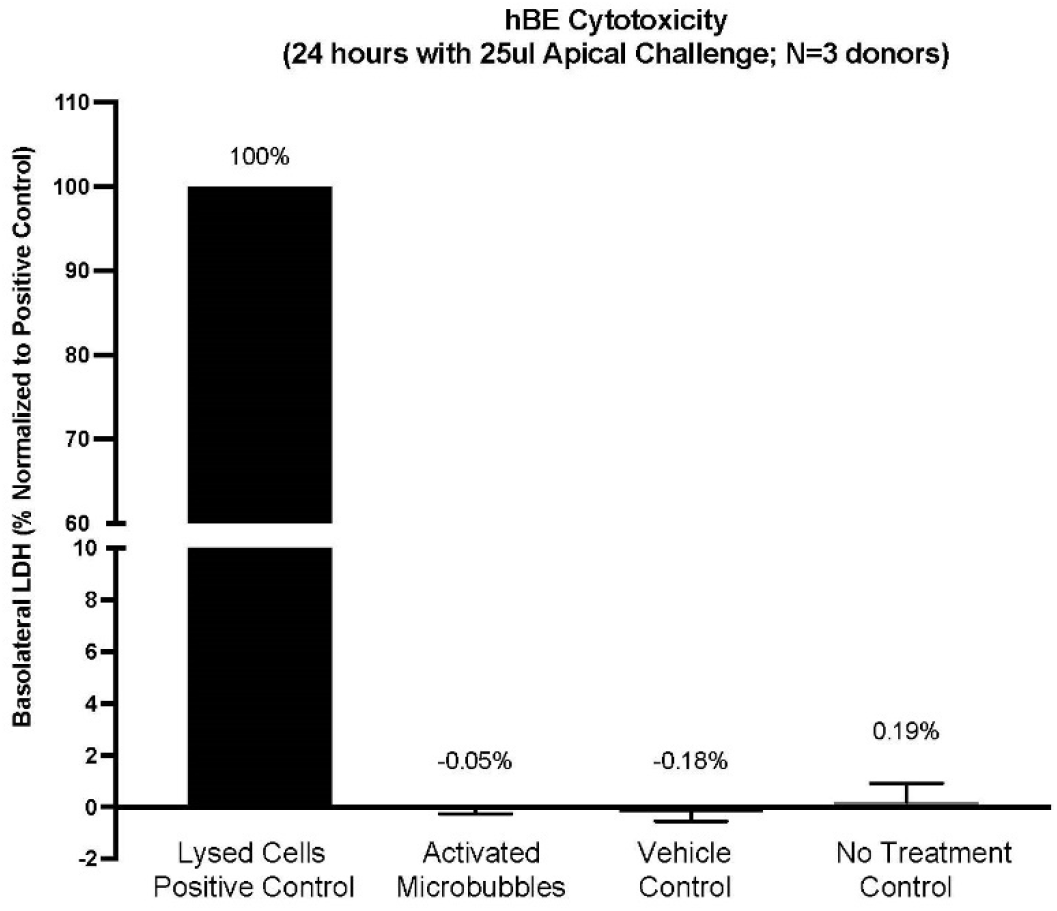
Mean (n=3 donor cultures) lactate dehydrogenase (LDH) release following exposure to microbubble contrast agent, vehicle control, or no treatment control. Basolateral LDH content in each condition was normalized to the positive control condition (maximal LDH release).

### Inflammatory response of MCA to cultured primary hBE cells

The basolateral cell culture medium was assessed to determine the presence of the proinflammatory cytokine interleukin-8 (IL-8) following 24-hour apical exposure to 25 μL of undiluted activated MCA, 25 μL of undiluted vehicle control, or a no treatment control. Neither exposure condition tested produced significant increases in basolateral IL-8 secretion as compared to the no treatment control (**Figure 5**).

**Figure 5:**
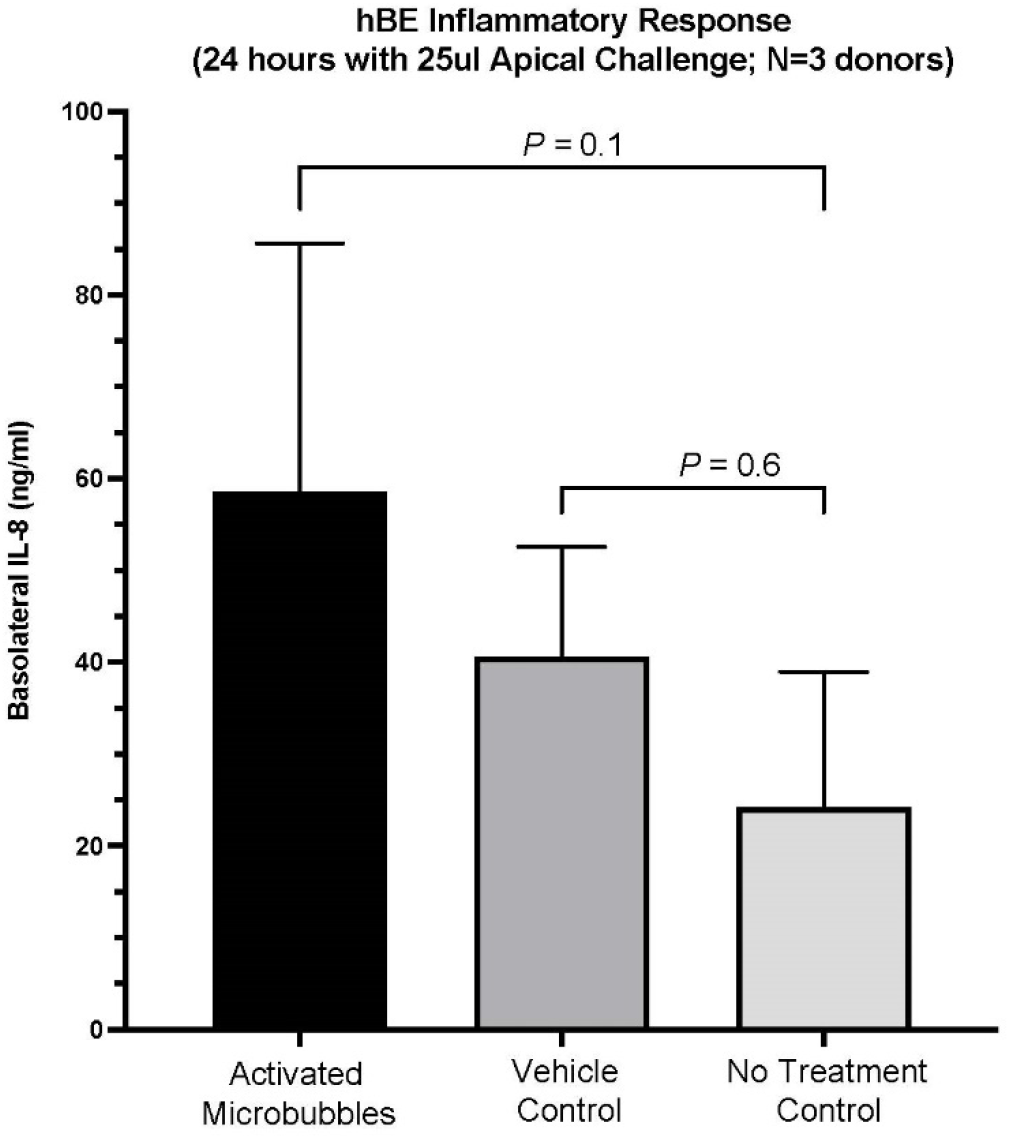
Mean (n=3 donor cultures) interleukin-8 (IL-8) content in basolateral medium following 24-hour exposure to activated microbubble contrast agent (MCA), nonactivated MCA, or no treatment (control). Means were compared with no treatment control using one-way ANOVA and Tukey’s multiple comparison test

## Discussion

In this feasibility study, we demonstrate the following: 1) Administration of aerosolized (nebulized) MCA through an *ex vivo* porcine trachea results in effective echo brightening and visualization of the tracheal wall contours, 2) 24-hour undiluted MCA challenge has no significant cytotoxic effects on cultured primary hBE cells, and 3) MCA does not illicit significant hBE cell proinflammatory responses after a 24-hour apical exposure.

An inhalable MCA may have several potential applications in point-of-care settings. One example would be providing visual guidance for tracheal intubation. Tracheal intubation is indicated for the management of acute respiratory distress syndromes (*e*.*g*., severe COVID-19 infection),^6^ and is also routinely performed for surgical procedures involving general anesthesia. Intubation is complicated by “difficult airways”, or anatomic distortions of the upper respiratory tract or trachea.^7^ Difficult tracheal intubation occurs most frequently in critically ill patients, with first-attempt failure rates as high as 30%.^8^ There is evidence to suggest that rates of both difficult intubations (requiring more than 3 attempts) and failed intubations (abandoned attempts, resorting to tracheostomy) have declined with standardized treatment algorithms.^9^ Nonetheless, the clinical sequelae of failed intubation can be severe, including brain damage and death.^10^ The incidence of failed intubation for pregnant women undergoing obstetric general anesthesia (2.6 per 1,000 anesthetics) has not changed since 1970,^11^ and outcomes of difficult intubation remain poor for all patients.^12^ Video laryngoscopes facilitate intubation by differentiating the trachea from esophagus, but are limited to imaging the upper airway and glottis. End-tidal capnography may be performed to confirm successful intubation into the trachea, but is unreliable in the setting of cardiac arrest, pulmonary edema, or pulmonary embolism.^13,14^ To assess endotracheal tube depth and confirm successful intubation, tracheal imaging by ultrasound, particularly point-of-care ultrasound (POCUS), has been increasingly utilized.^15^ However, the current imaging technique is suboptimal. Visualization can sometimes be improved by twisting the endotracheal tube from side-to-side.^16^ Other techniques, such as color Doppler or leaving the stylet in place to serve as a visual landmark, have not been shown effective.^17,18^ Here, we present proof-of-concept for an inhalable MCA which may facilitate ultrasound-guided tracheal tube placement and assessment of tracheal tube depth.

Another potential application for large airway evaluation by POCUS is detection of airway bleeds. Airway bleeds may arise from trauma, post-tonsillectomy hemorrhage, or other surgical etiologies, and are life-threatening even to young healthy patients due to potential aspiration of blood into the lungs.^19^ Current techniques to image airway bleeds, such as optical bronchoscopy, are particularly ineffective if blood from the oral or nasal cavity obscures the camera face as it is advanced into the trachea. When administered intravenously, MCAs strongly brighten sonographic echoes from the blood and opacify the blood-filled cardiac chambers in echocardiograms. With a respiratory route of administration, MCA may have the potential to rapidly enable sonographic visualization of airway bleeds in point-of-care settings.

Contrast-enhanced airway ultrasound may also be beneficial for monitoring pediatric patients with large airway disease. Cystic fibrosis (CF), a large airway disease, arises from mutations to the Cystic Fibrosis Transmembrane Conductance Regulator (*CFTR*) gene, and affects approximately 35,000 people in the United States.^20^ Once considered a disease of childhood, the life expectancy of individuals with CF has now increased to 48 years.^20^ The expanded life expectancy demands consideration of cumulative radiation exposure and its long-term effects. Across a 17 year period from 1992 – 2009, the mean cumulative effective dose (CED) of radiation exposure per patient with CF increased from 0.39 mSv/year to 1.67 mSv/year, with thoracic imaging accounting for 47% of the total exposure.^21^ A 6-fold increase in the use of computed tomography was also observed over this time interval.^21^ MDCT is not only used to monitor large airway pathology in patients with CF, but also response to therapies.^2^ Given the need for recurrent thoracic imaging, and the potential long-term effects of ionizing radiation, ultrasound may be a preferred alternative. Inhalable MCAs could have the potential to transform ultrasound into a powerful imaging modality for assessment and monitoring of CF and other large airway disease affecting pediatric patients.

In conclusion, we report promising sonographic enhancement and delineation of porcine *ex vivo* tracheal walls by nebulized MCA, with no significant cytotoxic or inflammatory effects on primary hBE cells. Our proof-of-concept study supports the feasibility of a respiratory route of MCA administration for clinical imaging. However, the safety of an inhalable MCA should be tested more extensively *in vitro* and *in vivo*, with preclinical evaluations conducted to assess acute lung tolerance, bioavailability, and clearance. The effectiveness of airway enhancement by nebulized MCA should also be evaluated, with radiologist-rated interpretations of image enhancement statistically analyzed in a well-powered study.

## Acknowledgments

The authors thank Dr. Prabir Roy-Chaudhury and Dr. Unimunkh Uriyanghai for providing an *ex vivo* porcine trachea for use in our experiments.

## Bibliography

1. Hansell DM, Bankier AA, MacMahon H, McLoud TC, Müller NL, Remy J. Fleischner Society: Glossary of Terms for Thoracic Imaging. Radiology 2008;246:697–722. Available from: http://pubs.rsna.org/doi/10.1148/radiol.2462070712

2. Semple T, Calder A, Owens CM, Padley S. Current and future approaches to large airways imaging in adults and children. Clin Radiol 2017;72:356–374. Available from: https://linkinghub.elsevier.com/retrieve/pii/S0009926017300405

3. Wei K, Mulvagh SL, Carson L, Davidoff R, Gabriel R, Grimm RA, Wilson S, Fane L, Herzog CA, Zoghbi WA, Taylor R, Farrar M, Chaudhry FA, Porter TR, Irani W, Lang RM. The Safety of Definity and Optison for Ultrasound Image Enhancement: A Retrospective Analysis of 78,383 Administered Contrast Doses. J Am Soc Echocardiogr 2008;21:1202–1206. Available from: https://linkinghub.elsevier.com/retrieve/pii/S0894731708004379

4. Chong WK, Papadopoulou V, Dayton PA. Imaging with ultrasound contrast agents: current status and future. Abdom Radiol 2018;43:762–772. Available from: http://link.springer.com/10.1007/s00261-018-1516-1

5. Chomas JE, Dayton P, Allen J, Morgan K, Ferrara KW. Mechanisms of contrast agent destruction. IEEE Trans Ultrason Ferroelectr Freq Control 2001;48:232–248. Available from: http://ieeexplore.ieee.org/document/896136/

6. Hur K, Price CPE, Gray EL, Gulati RK, Maksimoski M, Racette SD, Schneider AL, Khanwalkar AR. Factors Associated With Intubation and Prolonged Intubation in Hospitalized Patients With COVID-19. Otolaryngol Neck Surg 2020;163:170–178. Available from: http://journals.sagepub.com/doi/10.1177/0194599820929640

7. Higgs A, McGrath BA, Goddard C, Rangasami J, Suntharalingam G, Gale R, Cook TM. Guidelines for the management of tracheal intubation in critically ill adults. Br J Anaesth 2018;120:323–352. Available from: https://linkinghub.elsevier.com/retrieve/pii/S000709121754060X

8. Martin LD, Mhyre JM, Shanks AM, Tremper KK, Kheterpal S. 3,423 Emergency Tracheal Intubations at a University Hospital. Anesthesiology 2011;114:42–48. Available from: https://pubs.asahq.org/anesthesiology/article/114/1/42/10948/3-423-Emergency-Tracheal-Intubations-at-a

9. Schroeder RA, Pollard R, Dhakal I, Cooter M, Aronson S, Grichnik K, Buhrman W, Kertai MD, Mathew JP, Stafford-Smith M. Temporal Trends in Difficult and Failed Tracheal Intubation in a Regional Community Anesthetic Practice. Anesthesiology 2018;128:502–510. Available from: https://pubs.asahq.org/anesthesiology/article/128/3/502/18877/Temporal-Trends-in-Difficult-and-Failed-Tracheal

10. Cook TM, Woodall N, Harper J, Benger J. Major complications of airway management in the UK: results of the Fourth National Audit Project of the Royal College of Anaesthetists and the Difficult Airway Society. Part 2: intensive care and emergency departments †. Br J Anaesth 2011;106:632–642. Available from: https://linkinghub.elsevier.com/retrieve/pii/S0007091217332105

11. Kinsella SM, Winton AL, Mushambi MC, Ramaswamy K, Swales H, Quinn AC, Popat M. Failed tracheal intubation during obstetric general anaesthesia: a literature review. Int J Obstet Anesth 2015;24:356–374. Available from: https://linkinghub.elsevier.com/retrieve/pii/S0959289X15000916

12. Joffe AM, Aziz MF, Posner KL, Duggan L V., Mincer SL, Domino KB. Management of Difficult Tracheal Intubation. Anesthesiology 2019;131:818–829. Available from: https://pubs.asahq.org/anesthesiology/article/131/4/818/897/Management-of-Difficult-Tracheal-IntubationA

13. Li J. Capnography alone is imperfect for endotracheal tube placement confirmation during emergency intubation. J Emerg Med 2001;20:223–229. Available from: https://linkinghub.elsevier.com/retrieve/pii/S0736467900003188

14. Takeda T, Tanigawa K, Tanaka H, Hayashi Y, Goto E, Tanaka K. The assessment of three methods to verify tracheal tube placement in the emergency setting. Resuscitation 2003;56:153–157. Available from: https://linkinghub.elsevier.com/retrieve/pii/S0300957202003453

15. Gottlieb M, Holladay D, Burns KM, Nakitende D, Bailitz J. Ultrasound for airway management: An evidence-based review for the emergency clinician. Am J Emerg Med 2020;38:1007–1013. Available from: https://linkinghub.elsevier.com/retrieve/pii/S0735675719308162

16. Gottlieb M, Burns K, Holladay D, Chottiner M, Shah S, Gore SR. Impact of endotracheal tube twisting on the diagnostic accuracy of ultrasound for intubation confirmation. Am J Emerg Med 2020;38:1332–1334. Available from: https://linkinghub.elsevier.com/retrieve/pii/S0735675719306904

17. Göksu E, Sayraç V, Oktay C, Kartal M, Akcimen M. How stylet use can effect confirmation of endotracheal tube position using ultrasound. Am J Emerg Med 2010;28:32–36. Available from: https://linkinghub.elsevier.com/retrieve/pii/S0735675708006633

18. Gottlieb M, Holladay D, Serici A, Shah S, Nakitende D. Comparison of color flow with standard ultrasound for the detection of endotracheal intubation. Am J Emerg Med 2018;36:1166–1169. Available from: https://linkinghub.elsevier.com/retrieve/pii/S073567571730966X

19. Kristensen MS, McGuire B. The Bloody and Bleeding Airway In: Core Topics in Airway Management. Cambridge University Press; 2020. p. 282–289.Available from: https://www.cambridge.org/core/product/identifier/9781108303477%23CN-bp-32/type/book_part

20. Cystic Fibrosis Foundation Patient Registry. 2019 Annual Data Report 2019;Available from: https://cff.org/Research/Researcher-Resources/Patient-Registry/2019-Patient-Registry-Annual-Data-Report.pdf

21. O’Connell OJ, McWilliams S, McGarrigle A, O’Connor OJ, Shanahan F, Mullane D, Eustace J, Maher MM, Plant BJ. Radiologic Imaging in Cystic Fibrosis. Chest 2012;141:1575–1583. Available from: https://linkinghub.elsevier.com/retrieve/pii/S0012369212603481

